# SplitMEM: Graphical pan-genome analysis with suffix skips

**DOI:** 10.1101/003954

**Authors:** Shoshana Marcus, Hayan Lee, Michael C. Schatz

## Abstract

**Motivation:** With the rise of improved sequencing technologies, genomics is expanding from a single reference per species paradigm into a more comprehensive pan-genome approach with multiple individuals represented and analyzed together. One of the most sophisticated data structures for representing an entire population of genomes is a compressed de Bruijn graph. The graph structure can robustly represent simple SNPs to complex structural variations far beyond what can be done from linear sequences alone. As such there is a strong need to develop algorithms that can efficiently construct and analyze these graphs.

**Results:** In this paper we explore the deep topological relationships between the suffix tree and the compressed de Bruijn graph. We introduce a novel *O*(*n* log *n*) time and space algorithm called splitMEM, that directly constructs the compressed de Bruijn graph for a pan-genome of total length *n*. To achieve this time complexity, we augment the suffix tree with *suffix skips*, a new construct that allows us to traverse several suffix links in constant time, and use them to efficiently decompose maximal exact matches (MEMs) into the graph nodes. We demonstrate the utility of splitMEM by analyzing the pan-genomes of 9 strains of *Bacillus anthracis* and 9 strains of *Escherichia coli* to reveal the properties of their core genomes.

**Availability:** The source code and documentation are available open-source at http://splitmem.sourceforge.net

## 1 INTRODUCTION

### 1.1 Background

Genome sequencing has rapidly advanced in the past 20 years. The first free living organism was sequenced in 1995, and since then the number of genomes sequenced per year has been growing at an exponential rate (Liolios *et al*., 2006). Today, there are currently nearly twenty thousand genomes sequenced across the tree of life, including reference genomes for hundreds of eukaryotic and thousands of microbial species. Reference genomes play an important role in genomics as an exemplar sequence for a species, and have been extremely successful at enabling genome resequencing projects, gene discovery, and numerous other important applications. However, reference genomes also suffer in that they represent a single individual or a mosaic of individuals as a single linear sequence, making them an incomplete catalog of all of the known genes, variants, and other variable elements in a population. Especially in the case of structural and other large-scale variations, this creates an analysis gap when modeling the role of complex variations or gene flow in the population. For the human genome, for example, multiple auxiliary databases including dbSNP, dbVAR, DGV, and several others must be separately queried through several different interfaces to access the population-wide status of a variant (MacDonald *et al*., 2014).

The “reference-centric” approach in genomics has been established largely because of technological and budgetary concerns. Especially in the case of mammalian-sized genomes, it remains prohibitively expensive and technically challenging to assemble each sample into a complete genome *de novo*, making it substantially cheaper and more accessible to analyze a new sample relative to an established reference. However, for some species, especially medically or otherwise biologically important microbial genomes, multiple genomes of the same species are available. In the current version of NCBI GenBank, 296 of the 1471 bacterial species listed have at least two strains present, including 9 strains of *Bacillus anthracis* (the etiologic agent of anthrax), 62 strains of *Escherichia coli* (the most widely studied prokaryotic model organism) and 72 strains of *Chlamydia trachomatis* (a sexually transmitted human pathogen). This was done because the different genomes may have radically different properties or substantially different gene content despite being of the same species: most strains of *E. coli* are harmless, but some are highly pathogenic (Rasko *et al*., 2011b).

When multiple genomes of the same or closely related species are available, the “pan-genome” of the population can be constructed, and analyzed as a single comprehensive catalog of all of the sequences and variants in the population. Several techniques and data structures have been proposed for representing the pan-genome, i.e. (Rasko *et al*., 2008). The most basic is a linear concatenation of the reference genome plus any novel sequences found in the population appended to the end or stored in a separate database such as dbVAR. The result is a relatively simple linear sequence, but also loses much of the value of population-wide representation, necessitating auxiliary tables to record the status of the concatenated sequences. More significantly, a composite linear sequence may have ambiguity or loss in information of how the population variants relate to each other, especially at positions where the sequences of the individuals in the population diverge, i.e. branch-points between sequences shared among all the strains to any strain-specific sequences and back again.

A much more powerful representation of a pan-genome is to represent the collection of genomes in a graph: sequences that are shared or unique in the population can be represented as nodes, and edges can represent branch points between shared and strain-specific sequences (Figure 1). More specifically, the de Bruijn graph is a robust and widely used data structure in genomics for representing sequence relationships and for pan-genome analysis (Iqbal *et al*., 2012). In the case of a pan-genome, we can color the de Bruijn graph to record which of the input genome(s) contributed each node. This way the complete pan-genome will be represented in a compact graphical representation such that the shared/strain-specific status of any substring is immediately identifiable, along with the context of the flanking sequences. This strategy also enables powerful topological analysis of the pan-genome not possible from a linear representation.

**Fig. 1.**
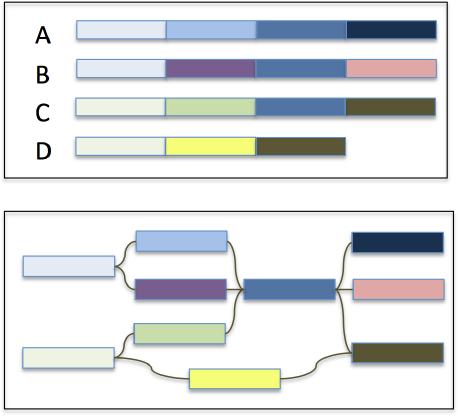
Overview of a graphical representation of a pan-genome. The four input genomes (A-D) are decomposed into segments shared or specific to the individuals in the population with edges maintaining the adjacencies of the segments.

As originally presented, the de Bruijn graph encodes each distinct length *k* substring as a node and includes a directed edge between substrings that overlap by *k* – 1 base pairs. However, many of the nodes and edges of a de Bruijn graph can be “compressed” whenever the path between two nodes is non-branching. Doing so often leads to a substantial savings in graph complexity and a more interpretable topology: in the case of a pan-genome graph, after compression nodes will represent variable length strings up to divergence in shared/strain-specific status or sequence divergence after a repeated sequence. The compressed de Bruijn graph is therefore the preferred data structure for pan-genome analysis, but it is not trivial to construct such a graph without first building the uncompressed graph, and then identifying and merging compressible edges, all of which requires substantial overhead. Here we present a novel space and time efficient algorithm called splitMEM for constructing the compressed de Bruijn graph from a generalized suffix tree of the input genomes. Our approach relies on the deep relationships between the topology of the suffix tree and the topology of the compressed de Bruijn graph, and leverages a novel construct we developed called *suffix skips* that makes it possible to rapidly navigate between overlapping suffixes in a suffix tree. We apply these techniques to study the pan-genomes of all 9 available strains of *B. anthracis* and a selection of 9 strains of *E. coli* to map and compare the “core genomes” of these populations. All of the source code and documentation for the analysis are available open-source at http://splitmem.sourceforge.net.

### 1.2 Problem Definition

The de Bruijn graph representation of a sequence contains a node for each distinct length *k* substring, called a *k*-mer. Two nodes are connected by a directed edge *u → v* for every instance where the *k*-mer represented by *v* occurs immediately after the *k*-mer represented by *u* at any position in the sequence. In other words, there is an edge if *u* occurs at position *i* and *v* occurs at position *i*+1. By construction, adjacent nodes will overlap by *k* – 1 characters, and the graph can include multiple edges connecting the same pair of nodes or self-loops representing overlapping tandem repeats. This definition of a de Bruijn graph differs from the traditional definition described in the mathematical literature that requires the graph to contain all length-*k* strings that can be formed from an alphabet rather than just those present in the sequence. The formulation of the de Bruijn graph used in this paper is commonly used in the sequence assembly literature, and we follow the same convention (Kingsford *et al*., 2010). Notably, the original genome sequence, before decomposing it into *k*-mers for the graph, corresponds to an Eulerian path through the de Bruijn graph visiting each edge exactly once. In the case of the pan-genome, we first concatenate the individual genomes together separated by a terminal character and discard any nodes or edges spanning the terminal character. The nodes are colored to indicate which genome(s) the node originated from so that each individual genome can be represented by a walk of nodes of consistent color.

A de Bruijn graph can be “compressed” by merging non-branching chains of nodes into a single node with a longer sequence. Suppose node *u* is the only predecessor of node *v* and *v* is the only successor of *u*. They can thus be unambiguously compressed without loss of sequence or topological information by merging the sequence of *u* with the sequence of *v* into a single node that has the predecessors of *u* and the successors of *v*. After maximally compressing the graph, every node will terminate at a “branch-point”, meaning every node has in-degree ≥ 2 or its single predecessor has out-degree ≥ 2 and every node has out-degree ≥ 2 or its single successor has in-degree ≥ 2. The compressed de Bruijn graph has the minimum number of nodes with which the path labels in the compressed graph are the same as in the uncompressed graph (Kingsford *et al*., 2010). In this way, the compressed de Bruijn graph of a pan-genome will naturally branch at the boundaries between sequences that diverge in their amount of sharing in the population.

The compressed de Bruijn graph is normally built from its uncompressed counterpart, necessitating the initial construction and storage of a much larger graph. In the limit, a basic construction algorithm may need to construct and compress *n* nodes while ours would directly output just a single node. In practice, the compressed graph of real genomic data is often orders of magnitude smaller than the uncompressed, although the exact savings is data dependent.

In this paper, we present an innovative algorithm that directly constructs the compressed de Bruijn graph by exploiting the relationships between the compressed de Bruijn graph and the suffix tree of the sequences. Our algorithm achieves overall *O*(*n* log *n*) time and space complexity for an input sequence of total length *n*. Alternatively, we also present a slower algorithm in the supplementary material that constructs the compressed de Bruijn graph from the set of exact self-alignments of length ≥ *k* in the genome. The alignment-based algorithm considers each alignment in turn and decomposes the graph nodes to represent smaller substrings when alignments are found to overlap one another. At worst, the number of pairwise alignments in a genome can be quadratic. Both algorithms have the same underlying intuition and the faster suffix-tree approach was inspired by the alignment-based algorithm.

### 1.3 Suffix Tree, Suffix Array, and MEMs

The suffix tree is a data structure that facilitates linear time solutions to many common problems in computational biology, such as genome alignment, finding the longest common substring among genomes, all-pairs suffix-prefix matching, and locating all maximal repetitions (Gusfield, 1997). It is a compact trie that represents all suffixes of the underlying text. The suffix tree for *T*=*t*_1_*t*_2_ … *t_n_* is a rooted, directed tree with *n* leaves, one for each suffix. A special character “$” is appended to the string before construction of the suffix tree to guarantee that each suffix ends at a leaf in the tree. Each internal node, except the root, has at least two children. Each edge is labeled with a nonempty substring of *T* and no two edges out of a node begin with the same character. The path from the root to leaf *i* spells suffix *T* [*i…n*].

The suffix tree can be constructed in linear time and space with respect to the string it represents (Ukkonen, 1995). Suffix links are an implementation technique that enable linear time and space suffix tree construction algorithms. Suffix links facilitate rapid navigation to a distant but related part of the tree. A suffix link is a pointer from an internal node representing a string *xS* to another internal node representing string *S*, where *x* is a single character and *S* is a possibly empty string.

A closely related data structure, called a *suffix array*, is an array of the integers in the range 1 to *n* specifying the lexicographic order of the *n* suffixes of string *T*. It can be obtained in linear time from the suffix tree for *T* by performing a depth-first traversal that traverses siblings in lexical order of their edge labels. (Gusfield, 1997) For any node *u* in the suffix tree, the subtree rooted at *u* contains one leaf for each suffix in a contiguous interval in the suffix array. That interval is the set of suffixes beginning with the path label from the root to node *u* (Kasai *et al*., 2001).

Maximal exact matches (MEMs) are exact matches within a sequence that cannot be extended to the left or right without introducing a mismatch. By construction, MEMs are internal nodes in the suffix tree that have *left-diverse* descendants, i.e., leaves that represent suffixes that have different characters preceding them in the sequence. As such, the MEM nodes can be identified in linear time by a bottom-up traversal of the tree, tracking the set of character preceding the leaves of the subtree rooted at each node. Since each MEM is an internal node in the suffix tree, there are at most *n* maximal repeats in a string of length *n* (Gusfield, 1997). Our algorithm computes the nodes in the compressed de Bruijn graph by decomposing the MEMs and extracting overlapping components that are of length ≥ *k*.

## 2 METHODS

In this section we describe our algorithm for constructing the compressed de Bruijn graph for a genome in *O*(*n* log *n*) time and space. It is outlined in Algorithm 1. The basis of our algorithm is deriving the set of compressed de Bruijn graph nodes from the set of *MEM_≥k_* nodes in the suffix tree, i.e., internal nodes that represent maximal exact matches of length ≥ *k* in the genome. The underlying algorithm was inspired by the use of the suffix tree to compute matching statistics as described by Gusfield (Gusfield, 1997).

Note that each node in the compressed de Bruijn graph is labeled by a maximal genomic substring of length ≥ *k* for which there are no internal overlaps, with the same or with a different genomic substring, of length ≥ *k*. As in the uncompressed counterpart, edges connect substrings that have a suffix-prefix match of length *k* – 1 in the genome. The nodes in the compressed de Bruijn graph fall into two categories: *uniqueNodes* represent a unique subsequence in the pan-genome and have a single start position; and *repeatNodes* represent subsequences that occur at least twice in the pan-genome, either as a repeat in a single genome or a segment shared by multiple genomes in the pan-genome population. *uniqueNodes* can be thought of as nodes that link between *repeatNodes*. As such, our graph construction algorithm begins by identifying the set of *repeatNodes*, from which it constructs the necessary edges and *uniqueNodes* along the way.

The set of *MEM_≥k_* and the *repeatNodes* represent the same subsequences of the genome, although there is not a one-to-one correspondence, especially in the case of overlapping or nested MEMs (Figure 2). A *MEM_≥k_* may need to be split into several *repeatNodes* when it has subsequences of length ≥ *k* in common with itself or another *MEM_≥k_*. Some *repeatNodes* are exactly MEMs in the genome, while other *repeatNodes* are parts of a MEM that lie between two embedded MEMs. Any subsequence of length ≥ *k* that is shared among MEMs is necessarily a MEM. Consequently, our algorithm processes the set of *MEM_≥k_* and split them into *repeatNodes* by extracting common subsequences of minimum length *k* among them. Whenever a MEM is split to remove a shared *repeatNode*, the split results in at least one MEM as a resulting segment and the other segment can be unique to this MEM.

**Fig. 2.**
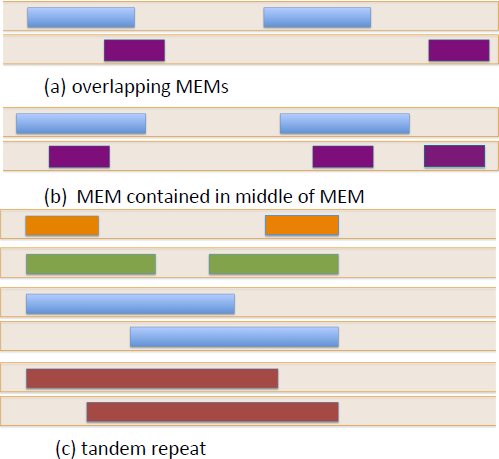
Different overlapping configurations of MEMs in a sequence. The colored blocks represent MEMs in a genomic sequence. Different colors are used for distinct MEMs.

### 2.1 Algorithm

The splitMEM algorithm uses a suffix tree of the genome to efficiently compute the set of *repeatNodes*. It builds a suffix tree of the pan-genome in linear time following Ukkonen’s algorithm (Ukkonen, 1995). It then marks internal nodes of the suffix tree that represent MEMs (or maximal repeats) of length ≥ *k*, in the suffix tree using linear time techniques of MUMmer (Kurtz *et al*., 2004) and preprocess the suffix tree for constant-time Lowest Marked Ancestor (LMA) queries in linear time. Then it constructs the set of *repeatNodes* by iterating through the set of *MEM_≥k_* in the suffix tree.

The challenge lies in identifying regions that are shared among *MEM_≥k_*s and decomposing *MEM_≥k_*s into the correct set of *repeatNodes*. If *m*_1_ and *m*_2_ are *MEM_≥k_*s and *m*_1_ occurs within *m*_2_, then *m*_1_ is a prefix of some suffix of *m*_2_. Thus, splitMEM can use the suffix links to iterate through the suffixes of *m*_2_ along with LMA queries to find the longest *MEM_≥k_*s that occurs at the beginning of each suffix. Each MEM is broken down to *repeatNodes* once and any embedded MEMs are extracted without examination. Thus, the subsequences that are shared among several MEMs are only decomposed once. We describe an efficient technique for constructing the set of *repeatNodes* in Section 2.2.

As an example, figure 3 shows the situation where a *MEM_≥k_* contains another *MEM_≥k_* within it. Two new *repeat nodes* are created for *xyzαβ*. One is the prefix ending after the first *k* – 1 characters of *α* (shown as ’ *α*) and the other is the suffix beginning with the last *k –* 1 characters of *α* (shown as *α*’). The smaller *MEM_≥k_* *α* is dealt with separately.

The positions at which the MEMs occur in the genome, and hence the start positions of the *repeatNodes*, can be quickly computed by considering the distance from the internal node to each leaf in its subtree and the genomic intervals that they represent. To make this computation efficient, we build a suffix array for the pan-genome and store at each suffix tree node its corresponding interval in the suffix array.

**Fig. 3.**
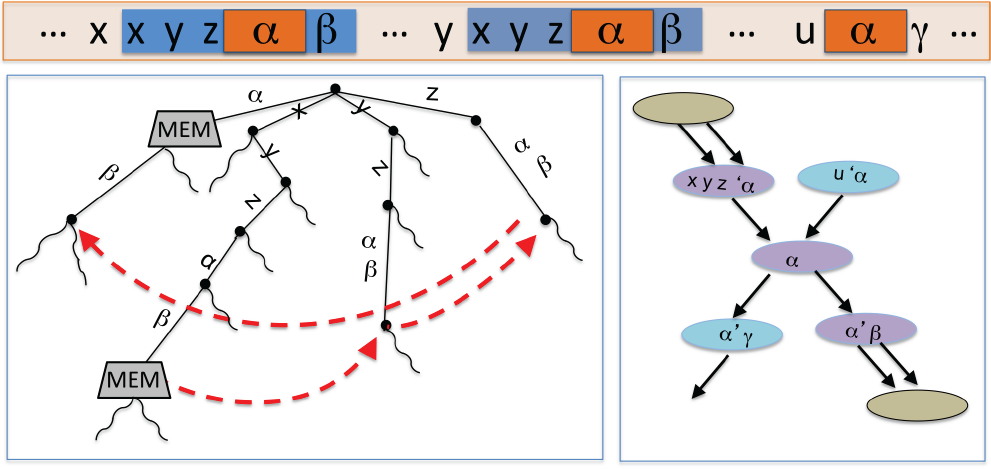
Part of the suffix tree for a genome (left) with the corresponding part of the compressed de Bruijn graph (right). Two MEMs in the suffix tree and the suffix links that are followed to decompose the larger MEM to at least three *repeat nodes*, the purple nodes in the graph on the right. x, y, and z are characters.α, β and γ are strings. Suffix links are displayed in red.

Once the algorithm has computed all the *repeatNodes*, it sorts the set of genomic starting positions that occur in each node so that it can construct the necessary set of edges between them in a single pass over this list. It also creates *uniqueNodes* to bridge any gaps between adjacent *repeatNodes* in the sorted list. It does this by iterating through the sorted list of start positions, *startPos* stored in each node. Suppose *startPos*[*i*] = *s*. It calculates the successive start position, *succ_i_*, from *s* and the length of the node containing *s*. If *succ_i_* is a start position of an existing node, it must be at position *i* + 1 in the sorted list, and cannot occur within a *repeatNode*. If *startPos*[*i* + 1] is a different value, the algorithm creates a *uniqueNode* to bridge the gap between startPos[*i*] and startPos[*i* + 1]. Then it creates an edge to join start position *s* to its successor, whether it’s in a *repeatNode* or a *uniqueNode*. If a *uniqueNode* was created, it also creates an edge to connect the new *uniqueNode* to its successor at *startPos*[*i* + 1].

The total length of all MEMs can be quadratic in the genome. Yet the total time complexity of Algorithm 1 is dependent on the total length of all repeat nodes, which is bounded by the genome size. Algorithm 1 runs in *O*(*n* log *n*) time and *O*(*n* + |*CDG*|) space, where |*CDG*| is the size of the compressed de Bruijn graph. We describe a technique in the next subsection that enables Algorithm 1 to achieve this time complexity.

### 2.2 Computing *repeatNodes* quickly with suffix skips

In this section we describe an *O*(*n* log *n*) time algorithm for deriving the set of *repeatNodes* from *MEM_≥k_*s in the suffix tree. It simulates the steps of iteratively traversing suffix links and performing an LMA query at each node traversed. In its basic form, as depicted in Figure 3, this process takes a total of *O*(*n*^2^) time, linear in the total length of all MEMs in the genome.

To accelerate it to *O*(*n* log *n*) time, we introduce *suffix skips* that generalize suffix links to trim more than a single character from the path to an internal node. Instead, suffix links trim *c* characters from the beginning of the path from the root to an internal node and navigate to the corresponding internal node in *O*(log *c*) time, instead of the *O*(*c*) time that would be required to traverse *c* suffix links (See Supplementary Figure 2). To compute the suffix skips, the algorithm creates a table of suffix skip pointers at each node *u*, with ⌊log_2_(*strdepth*(*u*))⌋ entries, where *strdepth*(*u*) is the length of the path from the root to node *u*. Entry *i* corresponds to the node that can be reached by traversing 2*^i^* suffix links from the node, *i* ≥ 0. The table is initialized with the original suffix link in entry 0 and then iteratively updated so that entry *i* of node *u* is assigned entry *i –* 1 of the node pointed to by node *u*’s *i –* 1th pointer: u → suffixSkip[*i*] = u suffixSkip[*i –* 1] → suffixSkip[*i –* 1]. Suffix skips are similar to the pointer jumping technique which speed up many parallel algorithms (Jaja, 1992).

Supplementary Algorithm 3 describes the use of *suffix skips* in an *O*(*n* log *n*) time procedure for deriving the *repeatNodes* from MEMs in the suffix tree. The algorithm iterates through the set of internal nodes that are marked as MEMs. For a MEM that is not a child of the root, we extend the node to include the path from the root to the internal node. The first LMA query identifies a potential prefix MEM. Then, embedded MEMs are identified by LMA queries and extracted by traversing *suffix skips*.A *repeatNode* is created to bridge gaps between embedded MEMs. If at any point a marked ancestor is found that extends to the end of the entire MEM, the process is complete. Otherwise, the last step is to create a *repeatNode* that spans the remaining suffix of the MEM.

We store auxiliary tables along with the *suffix skips* so that our algorithm can take advantage of suffix skips without potentially missing any nested MEMs. Along with each suffix skip stored at a node, we maintain a pointer to the bypassed LMA that is closest to the end of the destination node along with its base pair proximity to the end of the node. The speedup of *suffix skips* yields an algorithm with *O*(*n* log *n*) time complexity but requires an additional *O*(*n* log *n*) working space. To conserve space we only store suffix skips and auxiliary tables for nodes that can be traversed to decompose MEMs into *repeatNodes*, i.e., internal nodes that have string depth less than or equal to that of the longest *MEM_≥k_* and can be on the path of suffix links from a *MEM_≥k_* to the root.

### 2.3 Another speedup

We observe that a node is a MEM iff its ancestors are all MEMs. This allows us to save additional time when we decompose MEMs into *repeatNodes*. We use depth-first search to iterate over the suffix tree nodes to find *MEM_≥k_*s. Upon reaching a node *u* that has string depth ≥ *k* and is not a MEM, we bypass the subtree rooted at *u*.

## 3 RESULTS

We implemented Algorithm 1 along with Supplementary Algorithm 3 in C++ and made it available open-source as the splitMEM software. The code has been optimized for pan-genome and multi-*k*-mer analysis such that it can construct the graphs for several values of *k* iteratively without rebuilding the suffix tree. Using the software, we built compressed de Bruijn graphs for the pan-genomes of main chromosomes of two species: the 9 strains of *Bacillus anthracis* and a selection of 9 strains of *Escherichia coli* using the *k*-mer lengths 25, 100, and 1000bp (accessions listed in Supplementary Table 1). The three different *k*-mer lengths provide different contexts for localizing the graphs: shorter values provide higher resolution, while longer values will be more robust to repeats and link variations in close proximity into a single event. The overall characteristics of the pan-genome graphs are presented in Table 1 and renderings of the graphs are depicted in Supplementary Figures 3 - 8.

**Table 1.** *E. coli* and *B. anthracis* pan-genome graph characteristics.

| Species | K | Nodes | Edges | Avg. Degree |
| --- | --- | --- | --- | --- |
| B. anthracis | 25 | 103926 | 138468 | 1.33 |
| B. anthracis | 100 | 41343 | 54954 | 1.32 |
| B. anthracis | 1000 | 6627 | 8659 | 1.30 |
| E. coli | 25 | 494783 | 662081 | 1.33 |
| E. coli | 100 | 230996 | 308256 | 1.33 |
| E. coli | 1000 | 11900 | 15695 | 1.31 |

The pan-genome graphs of the two species show similar topologies, although for a given value of *k* the *E. coli* graph has 2 to 4 times as many nodes and edges than *B. anthracis*. In both cases, the node length distributions are exponentially distributed as shown in Supplementary Figure 9. For example, the mean node length for *B. anthracis* with *k*-mers of length 100 is 382bp and extending to as long as 451,679bp. The sharp peak at 199bp occurs from an enrichment of mutations where subpopulations or individual strains in the population differ by isolated single nucleotides more than *k* + 1 bp apart. At these sites, a “bubble” will form in the graph with a pair of nodes that are 2 * *k–*1 bp long capturing all of the *k*-mers that intersect the variant. Mutations of more than a single base form bubbles with nodes that are 2 * *k –* 1+ *v* bp long, where *v* is the length of the variant. Copy number and other structural variants lead to more complex graph topologies but are all encoded in the pan-genome graph.

Figure 4 shows the levels of population-wide genome sharing among the nodes of the compressed de Bruijn graphs of the pan genomes with varying *k*-mer lengths. The sharing in *B. anthracis* is much higher than in *E. coli* across the levels of genome sharing.

This follows naturally from the high diversity of *E. coli* strains (Rasko *et al*., 2008), while many of the available sequences of *B. anthracis* were closely related and sequenced to track the origin of the Amerithrax anthrax attacks (Rasko *et al*., 2011a).

**Fig. 4.**
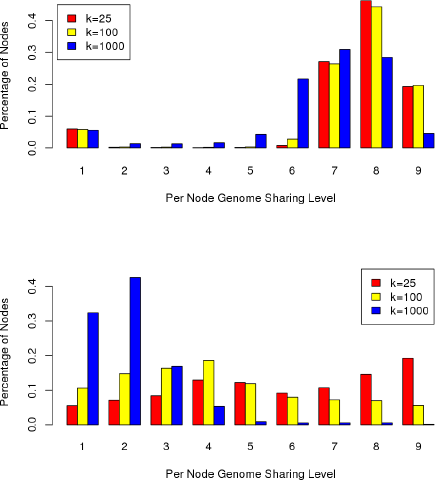
Levels of genome sharing in the nodes of the pan-genome graphs of 9 strains of *B. anthracis* (top) and *E. coli* (bottom). The plots show the fraction of nodes that have each level of sharing.

A major strength of a graphical pan-genome representation is that in addition to determining the shared or genome-specific sequences, the graph also encodes the sequence context of the different segments. We define the *core genome* to be the subsequences of the pan-genome that occur in at least 70% of the underlying genomes.

### Algorithm 1 Construct Compressed de Bruijn Graph from Suffix Tree

~~~
**Input:** genome sequence, *k*.
**Output:** compressed forward de Bruijn graph of genome.

**Compute set of *repeatNodes*.**
Build suffix tree of genome
Mark internal nodes in the suffix tree that represent MEMs of length ≥ k
Preprocess suffix tree for lowest marked ancestor (LMA) queries

**Split MEMs to *repeatNodes*.**
**for all** marked nodes *do*
                                                                       ► find k-mers shared with other MEMs or this MEM
 **while** node.strdepth ≥ *k* **do**
  **if** node has marked ancestor **then**
    create repeatNode to represent substring of MEM skipped by suffix link traversal since last internal MEM was removed
    follow suffix links to trim LMA from node
    continue traversing suffix links for any marked ancestors encountered during suffix link traversal, if they extend further
  **else**
    follow suffix link
  **end if**
 **end while**
 create *repeatNode* representing suffix of MEM that extends past last embedded MEM
**end for**

**Sort list of start positions in *repeatNodes***, with pointers to corresponding nodes.

**Compute outgoing edges for each node. Construct *uniqueNodes* along the way.**

**for all** startPos[*i*]=s **do**
  compute start position of successor *j*
  **if** startPos[*i* + 1]. *j* **then**
    create edge from node with s to node with *j*
  **else**
    create *uniqueNode* representing the subsequence from *j* until startPos[*i* + 1]
    create edge from node with *s* to node with *j*
  **end if**
**end for**
~~~

We computed the distance of each non-core node to the core genome in python using NetworkX with a branch-and-bound search intuited by Dijkstra’s algorithm for shortest path. Note a breadth-first search is not sufficient since two nodes can be further apart in terms of hops while they are actually closer neighbors with respect to base-pair distance along the path separating them. It traverses all distinct paths emanating from the source node until either a core node is reached or the current node was found to already have been visited by some shorter path. Once a path is found from the source node to the core genome, it uses this distance to bound the maximum search distances of the other candidate paths.

Using this approach, we performed both a forward search among descendants and a backwards search among predecessors to identify the distance to the closest core node and chose the minimum of these two distances in the two pan-genome graphs. This search takes *O*(*m*) time per source node, where *m* is the number of distinct edges in the graph. Thus, this computation takes a total runtime of *O*(*m * ℓ*) over all *ℓ* nodes in the graph. To keep the search tractable, we limited the search to a 1000-hop radius around each node. Supplementary Figure 10 shows the distribution of distances in the graphs. Overall, for *B.anthracis* most of the nodes were in the core genome since the strains are so similar or there was a very short path to the core genome. In contrast, the results for *E. coli* show the distribution of distances to the core genome follows an exponential distribution, suggesting a complex evolutionary history of mutations.

## 4 DISCUSSION

Comparative genomics has been and continues to be one of our most powerful tools for understanding the genome of a species. Now that genomic data are becoming more abundant, we are beginning to shift away from reference genomes and see the rise of pan-genomics. Already hundreds of microbial species have multiple complete genomes available, and through the rise of long read sequencing technologies from PacBio and other companies, we expect a rapid growth in the availability of complete genomes for additional bacterial and eukaryotic genomes (Koren *et al*., 2013; Roberts *et al*., 2013). Sequences that are highly conserved or segregating across the population can reveal clues about their phenotypic roles, and a comprehensive pan-genomic approach allows us to directly measure conservation in the context of the flanking sequences. The graphical pan-genome approach also consolidates all available information about complex structural variations and gene flow into a unified framework.

Our new *splitMEM* algorithm efficiently computes a graphical representation of the pan-genome by exploiting the deep relationships between suffix trees and compressed de Bruijn graphs. Maximal exact matches (MEMs) are readily identified in a suffix tree, and through the splitMEM algorithm are efficiently transformed into the nodes and edges of a compressed de Bruijn graph. This algorithm effectively unifies the most widely used sequence data structures in genomics into a single family containing suffix trees, suffix arrays, FM-indexes, and now compressed de Bruijn graphs. To accomplish this goal, we have proposed a new construct, called suffix skips, that generalizes the well-established concept of suffix links to interrelate more distantly related portions of the suffix tree.

To demonstrate the utility of the algorithm, we have applied it to analyze the pan-genomes of all 9 available *B. anthracis* genomes and a selection of 9 *E. coli* genomes. Interestingly, the distributions of the lengths of the nodes in the two pan-genome graphs are similar, other properties are markedly different, such as the distributions of the levels of sharing or the distance to the core genomes. This suggests that we have only narrowly explored the genomic variability of *B. anthracis* and future work remains to examine the functional significance of the variable regions.

Future work remains to improve splitMEM and further unify the family of sequence indices. Most desired are techniques that can leverage the compact space requirements of the FM-index for pan-genome analysis, along with techniques that can apply the suffix skip concept to generalize properties of the other indices, such as generalizing the Last-First property of the FM-index. We also aim to research additional downstream analysis algorithms for the pan-genome, especially developing a sequence aligner which can align directly to the graph structure. Finally, we also aim to extend the splitMEM algorithm to become more robust in the presence of incomplete genomes, so that fragmented draft genomes can be more readily analyzed.

## ACKNOWLEDGEMENTS

The authors would like to acknowledge Steven Skiena, Art Delcher, Adam Phillippy, Cole Trapnell, Mihai Pop, and Steven Salzberg for helpful discussions leading to this work. The project was supported in part by National Institutes of Health award R01-HG006677 and National Science Foundation award DBI-126383 to MCS.

